# Untangling or further entangling? Revelation of the complicated world of nucleic acid quadruplex folds

**DOI:** 10.64898/2025.12.14.694268

**Authors:** Thenmalarchelvi Rathinavelan, Sruthi Sundaresan

**Author notes:** Equal contribution.

## Abstract

Quadruplexes are four-stranded nucleic acid secondary structures playing diverse biological roles. A recent meticulous analysis of known quadruplex structures formed from synthetic oligonucleotides shows that quadruplexes can take 33 distinct structural motifs (pair of stacked quartets) that act as scaffolds for simple (multi-layered quartets) and complex (multi-motifs) quadruplex folds. These motifs differ from one another in terms of interstrand connectivity and orientation. The connectivity is primarily governed by diagonal loops that connect alternate strands, and lateral and propeller loops that connect adjacent strands. These loops are absent in the tetramer, wherein interstrand orientation alone dictates the motif type. Nevertheless, a key question that remains unresolved is the complete structural landscape of quadruplex motifs, limiting our understanding of the fold-dependency in various biological functions and diseases. To address this, a total of 156 unique theoretically possible quadruplex motifs are mathematically devised here. A tool named 3D-NuS-Qplex (https://project.iith.ac.in/3d-nus-qplex/) has also been developed to model all the theoretically possible energy-minimized quadruplex folds with the options to incorporate overhangs and non-G-quartets (in addition to G-quartets) in both DNA and RNA contexts. The diversity of quadruplex motifs reported here provides new insight into their structural complexity, which appears greater than previously anticipated.

**Highlights:**

- Quadruplex folds can be segmented into two-layered structural motifs that act as their scaffold
- 156 unique theoretically possible quadruplex motifs are mathematically devised, and an option to model them is discussed
- This reveals the complexity of quadruplex structures and offers new perspectives on quadruplex biology and disease

## Introduction

Nucleic acids take a wide range of secondary structures besides the right-handed double helical B-form [1-5]. One among them is a four-stranded nucleic acid structure known as a quadruplex or tetraplex, which was discovered even before the discovery of the double helical structure [6]. There are several evidences about the biological role of these secondary structures formed by DNA and RNA sequences [7-16]. Besides, these structures also play a crucial role in several disease conditions [17].

About 500 quadruplex structures, mostly corresponding to synthetic oligonucleotides, have been solved predominantly using X-ray crystallography followed by NMR [18, 19]. These structures are formed by G-rich sequences, which form G-quartets, wherein four guanines interact with each other through Hoogsteen hydrogen bonds (**Figure 1A, 1B**). Besides, nine non-G-quartets are also found to co-exist along with the G-quartets in a quadruplex structure (**Figure S1**).

**Figure 1.**
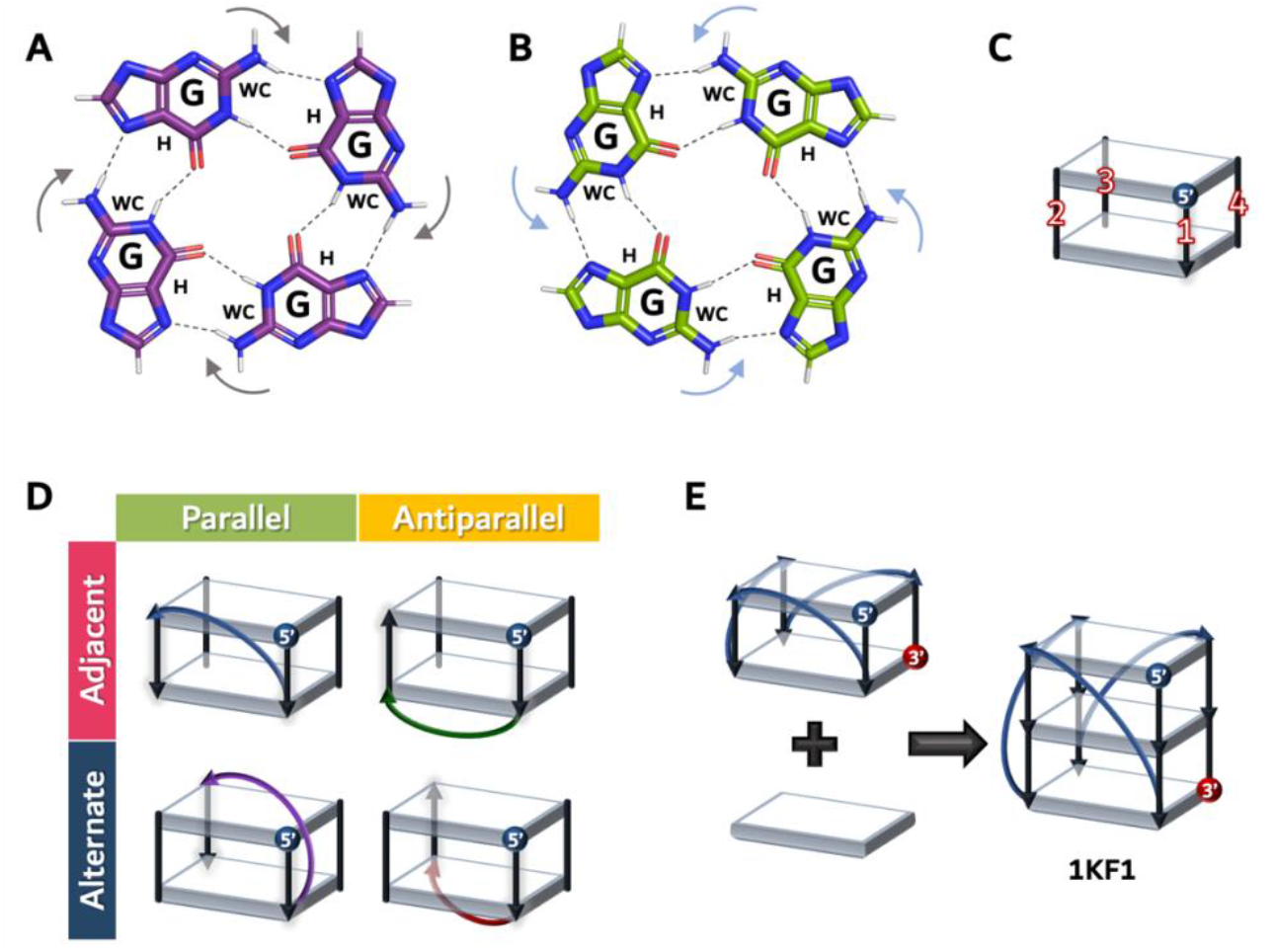
A schematic illustration of A) clockwise and B) anticlockwise Hoogsteen hydrogen bonding pattern (indicated by arrows) in a G-quartet. Hydrogen bonds are indicated in the dotted lines. “WC” and “H” represent Watson-Crick and Hoogsteen edges respectively. Note that the orientation of WC and H edges dictates the hydrogen bonding pattern. For example, when the WC edge of the bottom-right G is on the left side, it leads to the formation of a clockwise quartet (Left in A). One the other hand, if it is on the right side, leads to an anticlockwise quartet (Right in B). C) A representative illustration of a two layered quartet motif which acts as a building block for a fold. The number nomenclature marked here represents the strand numbers (1 to 4) of a quadruplex motif/fold and is followed throughout this article. The first strand always starts from the front-right with the 5’-end of the strand at the top. Note that the 5’ (blue colored circle) and 3’ (arrow head) ends of the 1^st^ strand alone is indicated for the illustration purpose since the directionality of the other strands varies depending on the motif type [18]. D) The definition of propeller (colored blue), lateral (colored green), diagonal-parallel (colored purple) and diagonal (colored red) loops connecting the different quadruplex strands are shown. E) An example illustrating how a motif can lead to the formation of a simple quadruplex fold [18].

There are reviews on different aspects of quadruplex folding patterns from time to time [18-22]. An earlier study has shown that G-rich sequences can fold into 26 monomers, considering the permissible loop combinations [21]. The same study revealed 16 possible *glycosyl* conformations for the G-quartets of the quadruplex fold. A recent article has shown, based on the analysis of experimentally determined structures, that quadruplexes can be represented in a modular fashion wherein a pair of stacked quartets, namely, motif (**Figure 1C**), serves as the scaffold for quadruplex folds. These structural motifs can either be a monomer (formed by a single sequence), dimer (2 sequences), trimer (3 sequences), or tetramer (4 sequences). It is noteworthy that each of these motifs has 4 strands (refer to **Figure 1C** for the definition of strand numbers). The same study identified 33 different motifs through the analysis of quadruplex structures deposited in PDB, which differ among themselves by the relative orientations between the four strands and the loops that connect them [18, 19]. Notably, if a loop connects the adjacent strands that interact through base…base hydrogen bonds in a parallel and antiparallel orientation, it is called a propeller and lateral loop, respectively (**Figure 1D**). Further, the diagonal loop connects two alternate strands (which are not involved in base…base hydrogen bonding interaction) in an antiparallel orientation (**Figure 1D**). Although a loop can theoretically connect two alternate strands (not involved in base…base hydrogen bonding interaction) in a parallel orientation (**Figure 1D**, henceforth referred to as diagonal-parallel), it has not yet been observed in the experimental structures. Another point to note is that a motif can even be discontinuous, *viz*., the adjacent quartets of all four strands in a quadruplex fold need not be connected [18]. A quadruplex fold can be formed using a single motif (**Figure 1E**), and a higher-order structure can be formed by combinations of motifs [18]. This further gives a clue that many undiscovered quadruplex structures may exist in nature, arising out of the combination of different motifs.

There are several attempts to develop algorithms (including machine learning) to predict the quadruplex folds from a given sequence [23-26]. Nonetheless, predicting the quadruplex structures from a given sequence remains a major challenge due to the lack of precise understanding of interstrand strand orientation and connectivity, which contribute to the diversity among quadruplex motifs. Additionally, structural diversity comes from a wide range of combinations that these motifs can adopt. Deriving the theoretically possible quadruplex motifs (the scaffold for the quadruplex folds), thus, would be highly instrumental in understanding the fold-dependent biological roles of quadruplexes. Thus, this study explores all theoretically possible continuous monomeric, dimeric, trimeric, and tetrameric quadruplex motifs. Further, a tool, namely, 3D-NuS-Qplex, has been developed to model all the theoretically derived quadruplex folds.

## Results and Discussion

### Theoretical formulation of possible quadruplex motifs

**Figure 2A** depicts the possible sequential arrangements among the four strands of a quadruplex motif. The encircled “1” (innermost circle) in **Figure 2A** represents the strand number 1 (which is the first strand) of the quadruplex motif (refer to **Figure 1C**), which progresses towards the outermost circle (which represents the last strand). The blue colored encircled number (**Figure 2A**) indicates the next strand (which can be strand number 2, 3, or 4 as shown in **Figure 1C**) that is connected to the 1^st^ strand of the quadruplex motif. Similarly, the next two consecutive strand numbers indicate the sequential order of the third (encircled in pink in **Figure 2A**) and fourth (encircled in green in **Figure 2A**) strands of the quadruplex motif, respectively. For instance, if the connectivity is between strand number 1 (first strand) and 4 (second strand) (triangle shaded in yellow color, **Figure 2A**), the next connectivity could be either to strand number 2 or 3 (**Figure 2A**). If the strand number 4 is connected to the strand number 3 (third strand), then the last strand (outermost circle, colored in green) of the quadruplex would be the strand number 2 (fourth strand). Thus, the strand connectivity goes in the following order: 1→4→3→2 in this example (triangle shaded in yellow color in **Figure 2A**). On the other hand, if the strand number 4 is connected to the strand number 2, it results in a motif whose connectivity is as follows: 1→4→2→3. Thus, four different quadruplex strands can be connected in six unique ways: 1→2→3→4; 1→2→4→3; 1→3→2→4; 1→3→4→2; 1→4→2→3; 1→4→3→2) (**Figure 2A**).

**Figure 2.**
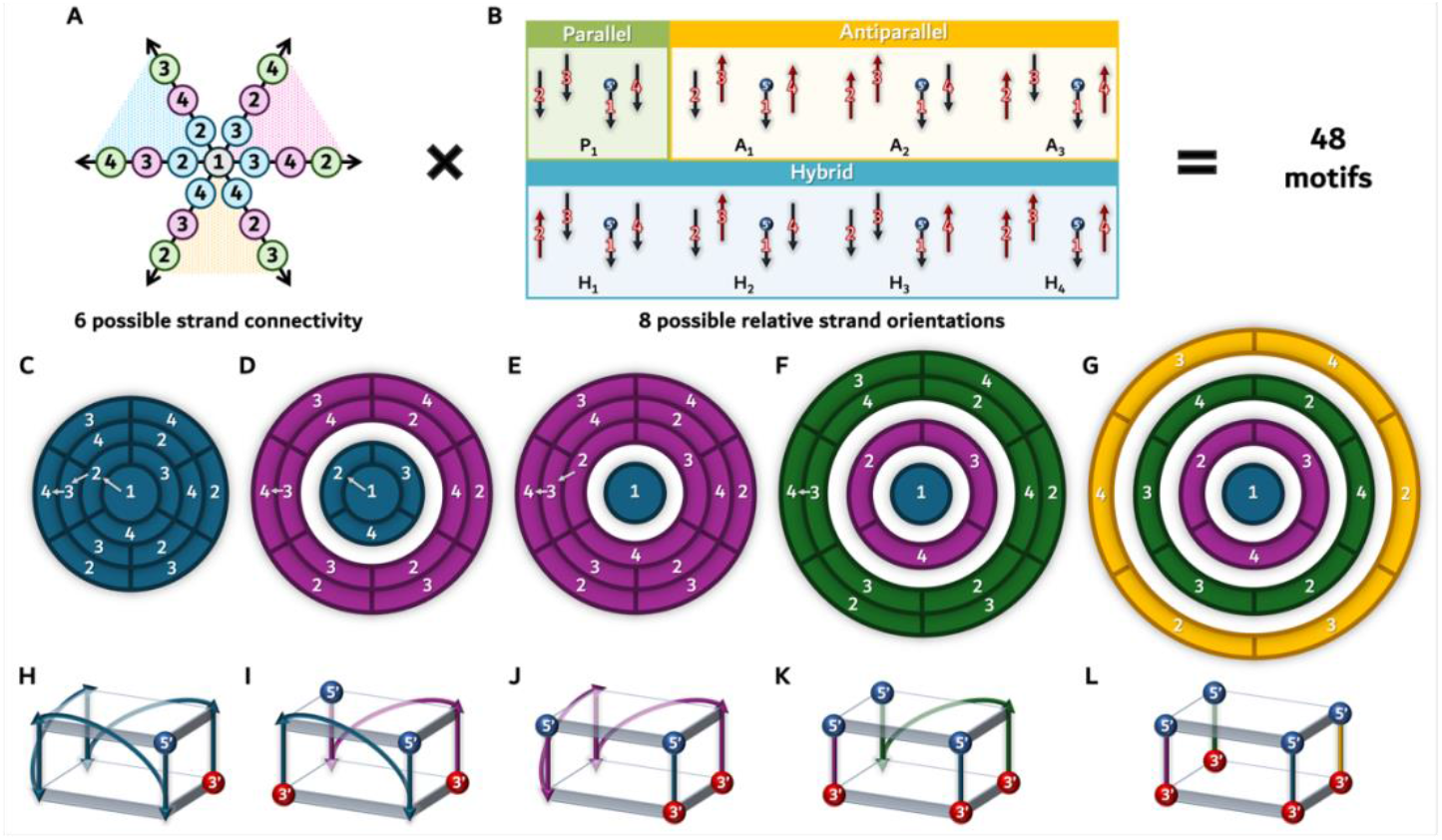
Diagram illustrating the theoretical formulation of possible quadruplex motifs. A) Six possible connectivity in quadruplex motifs. Note that the encircled numbers represent strand 1 to 4 of the quadruplex motif (**Figure 1C**). Note grey, blue, pink and green colored circles indicate the first, second, third and fourth strands of the motif in terms of the sequential arrangements among the quadruplex strands. B) Eight possible interstrand orientations in quadruplex motifs. The parallel (green), antiparallel (yellow) and hybrid (blue) motifs are indicated. Note that the tail and head of the arrows indicate 5’ and 3’ directions respectively. The number 48 indicates the total number of possible quadruplex motifs with the incorporation of 6 possible connectivity and 8 possible strand orientations. C) Monomeric motifs formed by a single sequence whose strands are connected by 3 loops. Note that each circle represent a quadruplex strand with the strand number marked. The strand numbers are indicated as per their sequential order in terms of connectivity with the central circle always being the strand number 1. Note that, the monomeric motif is uniformly represented by 4 blue colored circles as it is formed by a single sequence. D) Dimeric motifs that are made out of 2 sequences (colored in blue and purple) with each containing a loop. The first and second sequences of the dimeric fold are represented by blue and purple color respectively. E) Dimeric motifs formed by 2 sequences, where three strands of the motif are formed by a single sequence and are connected by 2 loops, while the other strand is independently formed by another sequence. The first and second sequences of the dimeric folds are represented by blue and purple color, respectively. F) Trimeric motifs made out of 3 sequences, in which, one sequence forms the two strands of the motif connected by a loop, while the other two sequences independently form the rest of the strands. The first, second and third sequences of the trimeric fold are represented by blue, purple and green color, respectively. G) Tetrameric motifs that are made out of 4 sequences (colored in blue, purple, green, and yellow circles), each forming 4 independent quadruplex strands (do not have any loop). Note that the filled circles in (C-G) represent the strands formed by a single sequence (wherein, the strands are connected by loop(s)), and the white space indicates (D-G) two independent sequences (*viz*., the two strands are not connected). (H-L) Representative quadruplex motifs corresponding to the monomer having 1→2→3→4 connectivity (H), and the concomitant dimers (I & J), trimer (K) and tetramer (L), whose interstrand connectivity is indicated by arrows in (C-G). Note that in (H-L) the first strand always starts from front-right with the 5’end on the top.

While the above description provides six possible connectivities between the different strand numbers of the quadruplex motif, incorporation of the strand directionality (*viz*., parallel[p] or antiparallel[a] orientation) brings further diversity to the quadruplex motif. For instance, in the above-given 1→4→3→2 motif, the strand number 1 can be connected to the strand number 4 in a parallel or an antiparallel fashion. Next, the strands numbers 4 and 3 can be parallel or antiparallel with respect to each other. Finally, the strands numbers 3 and 2 can also be connected in a parallel or an antiparallel orientation. Note that the orientation between the first and last strand is automatically taken care of. Considering these, 1→4→3→2 connectivity can lead to 8 different possible motifs by the inclusion of interstrand orientations: 1p→4p→3p→2, 1p→4p→3a→2, 1p→4a→3p→2, 1a→4p→3p→2, 1p→4a→3a→2, 1a→4p→3a→2, 1a→4a→3p→2, and 1a→4a→3a→2 (**Figure 2B)**. The eight orientations include, 1 parallel (all the 4 strands are in the same direction, denoted as P1 in **Figure 2B**), 3 antiparallel (2 strands are in one direction and 2 strands are in the opposite direction, denoted as A1-A3) and 4 hybrid (3 strands are in one direction and 1 strand is in the opposite direction, denoted as H1-H4) motifs respectively. Thus, incorporation of the strand directionality leads to eight different motifs (**Figure 2B**) for each of the 6 connectivities (**Figure 2A**). Thus, six different types of connectivity and 8 different interstrand directionality give rise to 48 quadruplex motifs (6 strand connectivity x 8 strand orientations = 48) (**Figure 2B, Table S1**).

When each of the aforementioned 48 motifs is formed by a single sequence, in which all 4 strands are continuously connected through three loops, it is called a monomeric motif. Note that the type of loops (propeller, lateral, diagonal, and diagonal-parallel, refer to **Figure 1D**) depends on the orientations of the connected strands. Considering this point, the diversity of the only possible parallel motif (P1 in **Figure 1B**, wherein all the strands are in parallel orientation) therefore arises out of the order of interstrand connectivity (**Figure 2A**) established through propeller and diagonal-parallel loops, as only they can connect 2 parallel strands. In contrast, lateral and diagonal loops do not come into the picture in the parallel motif as they can only connect antiparallel strands. Thus, based on the order of connectivity, 6 parallel motifs are possible. In addition to this, 18 antiparallel (denoted as A_1_, A_2_, and A_3_ in **Figure 2B**) motifs are possible when strand connectivity comes into the picture. Notably, in the cases of A_1_ and A_2_, starting from strand number 1, the interstrand orientations between the adjacent strands are alternately parallel and antiparallel. In A_1_, the strand numbers 1&2, 2&3, 3&4, and 4&1 are parallel, antiparallel, parallel, and antiparallel, respectively. In contrast, in the case of A_2_, the strand numbers 1&2, 2&3, 3&4, and 4&1 are antiparallel, parallel, antiparallel, and parallel, respectively. In A_3_, all the strands are antiparallel with respect to each other. Thus, there are 3 different classes of antiparallel motifs. Furthermore, each of these three motifs can lead to 6 different motifs based on the order of strand connectivity, as depicted in **Figure 2A**. In a similar fashion, each of the four different hybrid motifs (denoted as H_1_, H_2_, H_3_, and H_4_ in **Figure 2B**) leads to a total of 24 motifs. **Figure 2C** illustrates a different representation of **Figure 2A**, with the number indicated at each layer of the circle representing the sequential arrangement of connectivity between different quadruplex strand numbers.

Similarly, one can derive the number of possible dimeric motifs. Dimeric motifs can be derived from each monomeric motif by removing one loop from each of the 48 motifs. This can be possible in two different ways. In the first type, the dimer is formed by two independent sequences (indicated by the blue and purple colored circles in **Figure 2D**), with one loop in each sequence connecting two quadruplex strands. The second type of the dimeric fold is when the first strand of the quadruplex is formed by an independent sequence (indicated by the blue colored circle in **Figure 2E**) (Note that one can also keep any one of the other strands as the first strand which will essentially lead to the similar motifs.), whilst the remaining three strands of the quadruplex are formed by the second sequence (indicated by the purple colored circle in **Figure 2E**). Note that the white colored circle in the midst of the blue and purple circles (**Figure 2C-D**) indicates that the two strands are independent and not connected. As mentioned in the case of monomers, the direction of the sequence (5’ to 3’) goes from inner to outer circle (both in purple and blue colored circles). The aforementioned dimeric types can lead to 96 theoretical dimeric motifs (48 possible motifs in each type, based on strand orientation and connectivity, as discussed in the case of monomers).

Trimeric motifs can also be derived from monomeric motifs by removing two loops, or from dimeric motifs by removing one loop. The trimeric motif pattern is represented in **Figure 2F**, where the first strand is the innermost circle (indicated by a blue-colored circle). The second independent strand is shown in a purple circle (**Figure 2F**). The remaining two connected strands of the quadruplex are shown in green (**Figure 2F**). The 5’ to 3’ direction goes from the innermost blue colored circle to the outermost circle. Note that the white circle indicates that the two strands are independent and not connected. A total of 48 trimeric folds are possible based on strand orientation and connectivity.

Similarly, the tetrameric motifs can be derived from monomer, dimer, or trimer by the removal of three, two, and one loops, respectively. As shown in **Figure 2G**, the four strands (colored in blue, purple, green, and yellow) are not connected and independent (indicated by a white circle in).

**Figures 2H-2L** represent the monomeric 1→4→3→2 motif (**Figure 2H**) equivalent of dimeric (**Figure 2I, 2J**), trimer (**Figure 2K**), and tetrameric (**Figure 2L**) motifs.

### Theoretically possible unique quadruplex motifs

#### Monomeric motifs

Figure 3 illustrates all possible 48 monomers obtained from 6 interstrand connectivity (**Figure 2A**) and 8 interstrand orientations (**Figure 2B**). As discussed previously, there are 8 different orientations for each possible connectivity. In other words, there are 6 different connectivities for each possible strand orientation. In **Figure 3** under “1-2-3-4” column, the sequential order of connectivity between quadruplex motif strands are 1, 2, 3 and 4 which can have 8 different interstrand orientations: 1p-2p-3p-4 (MP1), 1p-2a-3p-4 (MA1), 1a-2p-3a-4 (MA7), 1a-2a-3a-4 (MA13), 1a-2a-3p-4 (MH1), 1p-2a-3a-4 (MH7), 1p-2p-3a-4 (MH13) and 1a-2p-3p-4 (MH19). Thus, a total of 6x8=48 monomeric (M) quadruplex motifs are possible based on the interstrand connectivity and orientations. Notably, according to the interstrand orientations, the connecting loop is decided as discussed previously. The parallel motif P_1_ (**Figure 2B**) has all four strands parallel to each other. However, the difference in interstrand connectivity among them leads to 6 different motifs (indicated in the first row of **Figure 3** under the “Parallel” label): 1p-2p-3p-4 (MP1), 1p-2p-4p-3 (MP2), 1p-3p-2p-4 (MP3), 1p-3p-4p-2 (MP4), 1p-4p-2p-3 (MP5) and 1p-4p-3p-2 (MP6), The MP1 motif (column1, row1) differs from MP2 (column 2, row1) due to the connectivity: while the former has the sequential order of strand connectivity as 1→2→3→4, the latter has 1→2→4→3. As discussed above, the adjacent parallel strands are connected through a propeller loop, while the alternate strands are connected through a diagonal-parallel loop (not observed in experimental structures) (**Figure 1D**). Yet another point is the crossing of diagonal-parallel loops observed in MP3 and MP4, which may be possible with a greater number of loop residues. While only MP1 and MP6 have been observed in experimental structures so far [18], one cannot deny the possibility of MP2, MP3, MP4, and MP5 motifs in nature.

**Figure 3.**
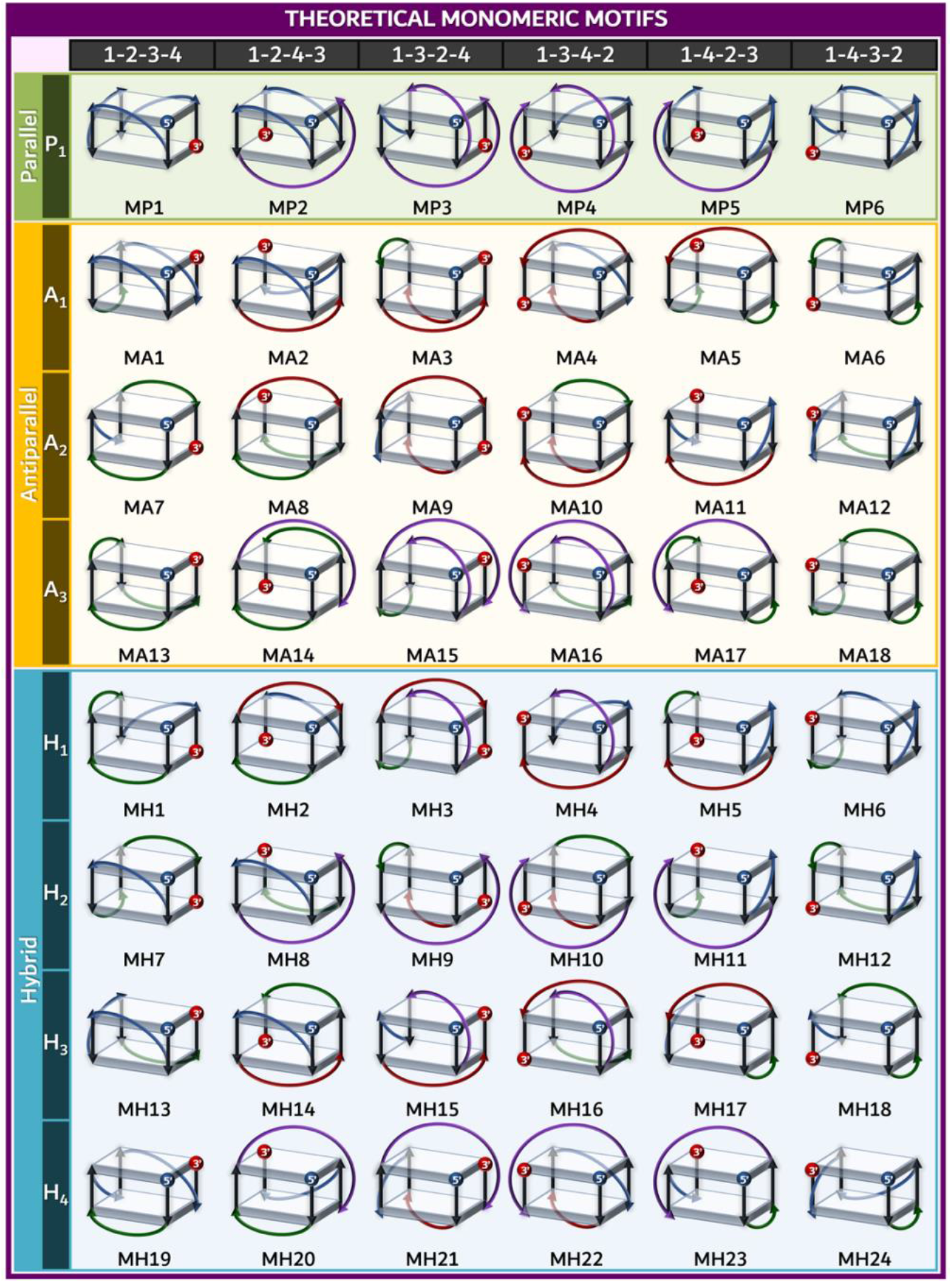
Theoretically possible 48 unique monomeric (M) quadruplex motifs derived out of 6 possible connectivity (column) and 8 possible orientations (row). “1-2-3-4”, “1-2-4-3”, “1-3-2-4”, “1-3-4-2”, “1-4-2-3” and “1-4-3-2” indicate 6 possible sequential orders of connectivity between different quadruplex strand numbers (**Figure 2A**). Note that “-” indicates that the sequence is continuous between different strands. 8 different interstrand orientations (**Figure 2B**) possible for each of the abovementioned connectivity is given under the respective rows. P_1_, A_1_-A_3_ and H_1_-H_4_ represent parallel, antiparallel and hybrid motif classes respectively (**Figure 2B**). The alphanumeric name corresponding to each motif is indicated adjacently. Note that a propeller loop that connects two adjacent strands parallelly is shown in blue, a lateral loop that connects two adjacent strands antiparallelly is in green, a diagonal loop that connects two alternate strands antiparallelly is in red, and a diagonal-parallel loop that connects alternate strands in a parallelly is in purple.

As explained in **Figure 2B**, there are 3 different classes (A_1_-A_3_) of antiparallel motifs. One can see from **Figure 3** that propeller, lateral, diagonal, and diagonal-parallel loops are all possible in antiparallel motifs; however, each motif includes a combination of at most two of these loop types. Notably, MP3 and MP4, MA3 and MA10 motifs (**Figure 3**) have 2 crossing diagonal loops, which may be sterically unfavorable [21] or require a greater number of loop residues. Hybrid motifs (H_1_-H_4_, **Figure 2B**) can have a combination of two or three types of loops (**Figure 3**). Thus, there are 48 unique monomeric motifs that do not share rotational symmetry with one another. This is because there is only one starting and ending position in a monomeric motif made up of a single sequence, and the starting position is always fixed to the top-right corner (**Figure 1C**).

#### Dimeric motifs

Typically, a dimeric (D) motif can be generated from a monomeric fold (MP1, **Figure 4A**) in three different ways. As shown in **Figure 4B**, removal of the propeller loop connecting strands 2 and 3 in the MP1 motif (**Figure 3**) leads to a dimeric motif (DP1) having 2 independent sequences, each with a propeller loop that connects 2 strands. On the other hand, the removal of the propeller between strand numbers 1 and 2 leads to a different type of dimeric motif (DP7) (**Figure 4C**), wherein 3 strands are connected by 2 consecutive propeller loops, while leaving the other strand independent. Although removal of the propeller loop between strand numbers 3 and 4 is possible (**Figure 4D**), it doesn’t lead to a different motif. This is because this motif is the same as the one arising from the removal of the propeller loop between strand numbers 1 and 2 (**Figure 4C**). This arises from the rotational symmetry shared by the two motifs (**Figure 4C,4D**). Thus, as discussed above, only 2 different dimeric motif patterns are possible from a monomer. This may theoretically result in 96 dimeric motifs (**Figure S2**) (instead of 144) arising out of each of the 48 monomeric motifs (**Figure 3**). Exceptionally, in some cases, dimeric motifs arising from two different monomeric motifs can eventually be the same. For instance, the removal of the diagonal-parallel loop between strand numbers 2 and 4 in MP2 (**Figure 3**) and strand numbers 4 and 2 in MP5 (**Figure 3**) leads to the same motif (DP2 and DP5 in **Figure S2**). This scenario arises from the motifs which are formed by two 2 independent sequences, each having a loop (**Figure S2**, shown above the dotted line). This reduces the 48 possible motifs to 32 (**Figure S2**). Thus, a total of 80 unique theoretical dimeric motifs are possible (**Figure S2**). As observed in the case of monomers, there is a possibility of two loops crossing over each other. This crossing can occur in different combinations, such as (diagonal)– (diagonal) (as in DA3, **Figure S2**), (diagonal-parallel)–(diagonal-parallel) (as in DA15, **Figure S2**), and (diagonal)–(diagonal-parallel) (as in DH3, **Figure S2**).

**Figure 4.**
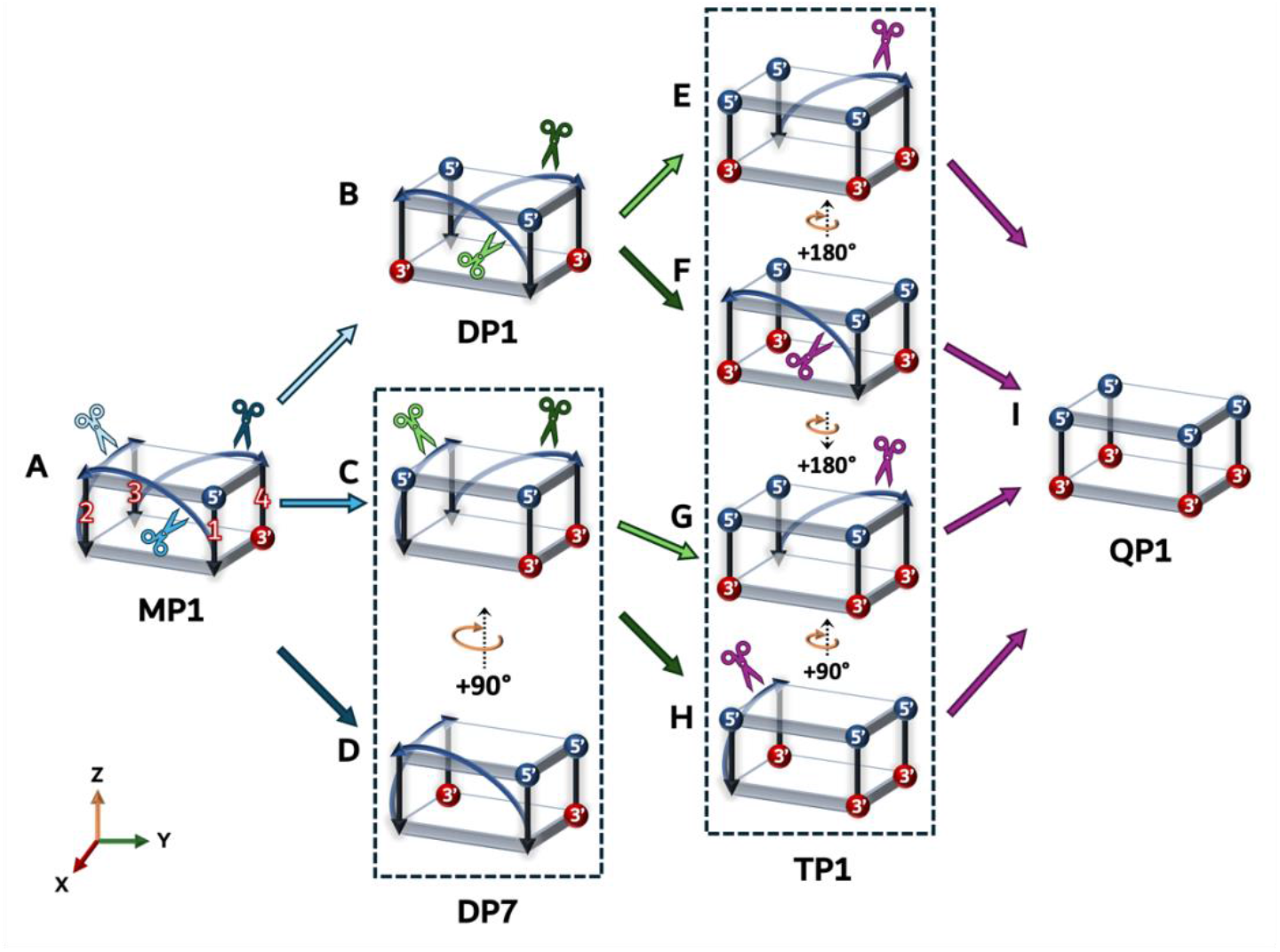
Derivation of dimeric (B-D), trimeric (E-H) and tetrameric (I) motifs from a monomeric motif by considering MP1 (A) as a case in point. While the 4 strands of the MP1 motifs are connected by three different loops, the other motifs can be derived by the removal of appropriate loops (represented by different colored scissors). Solid arrows are color coded according to the color of the scissor. Identical motifs are grouped under a dotted box. Note that circular arrows indicated along with the angles inside the dotted boxes represent the motifs that are related by a rotationally symmetry.

#### Trimeric motifs

Similar to the dimeric motif, a trimeric (T) motif can be derived from a dimeric motif by removing a loop, or from a monomeric motif by removing two loops. This leaves a motif with three different sequences, wherein only 2 strands are connected by a loop while the others stay unconnected and independent (**Figure 4E**). The example given in **Figure 4E** describes the derivation of a trimer motif from the dimer. Intriguingly, removal of any propeller loops in DP1 (**Figure 4B**) or DP7 (**Figure 4C and 4D**) leads to 4 possible motifs with 2 independent strands. Since any of these independent strands can be treated as the first strand, it leads to rotational symmetry; thus, there is only one unique trimeric motif. Thus, two different dimeric motifs, DP1 and DP7, derived from the same monomeric motif, MP1, again lead to one trimeric motif, TP1. Thus, one may envisage that there may be only 48 unique theoretical trimeric motifs (**Figure S3**). However, since only two strands are connected in a trimer, the trimers derived from two different monomeric/dimeric motifs can again lead to the same trimeric motif due to the rotational symmetry. One such example is MA1 (**Figure 3**) (leads to the DA1 and DA19 dimers, **Figure S2**) and MA11 (**Figure 3**) (leads to the DA11 and DA29 dimers, **Figure S2**) that lead to the same trimer TA1 (**Figure S3)**. Thus, the unique theoretical trimeric motifs reduce to 24 (**Figure S3**).

#### Tetrameric motifs

A tetrameric motif can be derived from a trimeric motif by removing the existing single loop (**Figure 4E-4H**). This results in four independent strands in the quadruplex motif. Thus, 6 connectivity doesn’t come into the picture, as any of the four strands (not connected and independent) can be the first strand. Thus, only 8 tetrameric motifs can be theoretically defined based on interstrand orientations (**Figure S4**). Furthermore, due to the rotational symmetry between two antiparallel motifs (A_1_ and A_2_ in **Figure 2B**), wherein 2 consecutive strands are in one direction, and the other 2 are in the opposite direction, both eventually lead to the same tetrameric motif. Similarly, all 4 hybrid motifs (H_1_-H_4_ in **Figure 2B**) result in a single motif. Thus, a total of 4 unique tetrameric motifs alone is possible.

Collectively, 156 unique quadruplex motifs are theoretically possible, which include 48, 80, 24, and 4 monomeric (**Figure 3**), dimeric (**Figure S2**), trimeric (**Figure S3**), and tetrameric (**Figure S4**) motifs, respectively. This is much higher than the so far known quadruplex motifs through experimental structures (33 motifs [18]). Perhaps this may be attributed to the influence of sequence and environmental conditions. This necessitates a detailed computational investigation, such as molecular dynamics simulations, to identify the energetically favourable motif(s) and their motif-sequence specificity. In any case, 156 unique quadruplex motifs reported here enlighten the importance of incorporating motif-sequence dependency in predicting quadruplex-forming regions using machine learning approaches.

## Limitations

The models generated can only be considered as an initial model and not a representation of the global minima. Further, complex quadruplex folds arising out of the combinations of different motifs (involving both left-handed and right-handed motifs) are not possible in the current implementation.

## Materials and Methods

### Implementation of a web interface for the automated modeling of quadruplex folds

The 200 motifs defined in **Figures 3, S2-S4** can be extended to a quadruplex fold, *viz*., a multi-layered G-quartet. This necessitates the need for the development of a tool that can model all the theoretically feasible quadruplex motifs to explore their biological relevance. Further, only a limited number of motifs are reported in experimental structures [18]. Indeed, an existing web tool models only a limited number of quadruplex folds [27]. Here, a web server, namely 3D-NuS-Qplex (https://project.iith.ac.in/3d-nus-qplex/), is developed to model these quadruplex folds by applying a helical twist (30°) and rise (3.4 Å) between the adjacent quartets. Note that a recent study has reported that the helical twist of quadruplex structures falls in the range of 10-40° [28]. Thus, to model all the theoretically possible quadruplex folds, 30° is chosen here as the helical twist. Similarly, the helical rise is chosen to be 3.4 Å. However, the values mentioned above are chosen to be constant across all the folds to model a broad range of quadruplex folds defined by all the motifs (**Figures 3, S2-S4**). Furthermore, 3D-NuS-Qplex also facilitates the use of nine different non-G-quartets (**Figure S1**), which are found alongside the G-quartets in experimentally determined structures deposited in the PDB [18, 19]. 3D-NuS-Qplex can also model quadruplex structures with 5’ and/or 3’ overhangs. The web interface of 3D-NuS-Qplex is shown in **Figure 5**.

**Figure 5:**
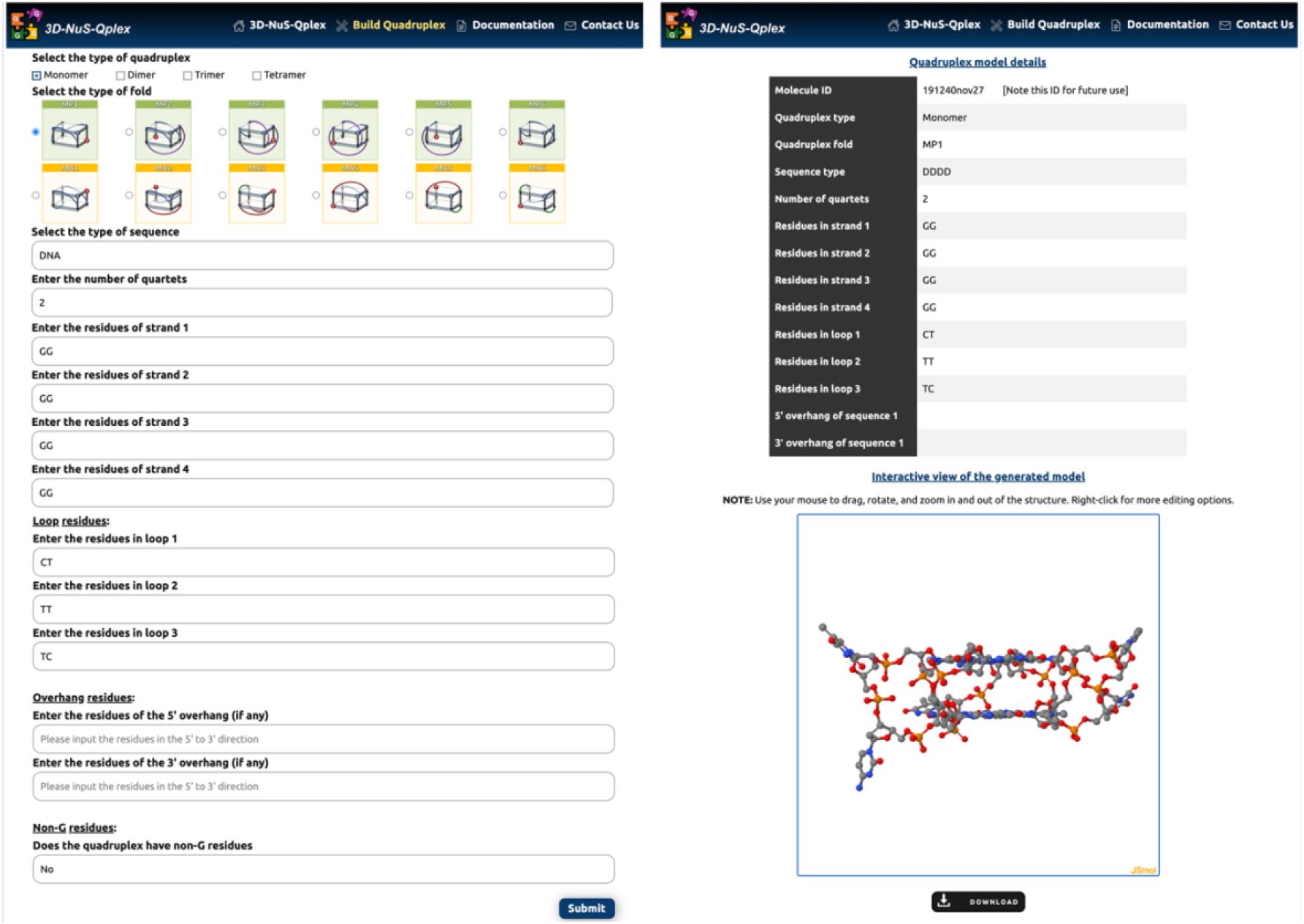
Snapshots of the 3D-NuS-Qplex interface. The workflow begins with the input interface shown on the left, where the user has to select the quadruplex type, fold type, the type of sequence (DNA/RNA), number of quartets, residues in each strand, loops, overhang residues (if any), and the positions of the non-G quartets (if any). After submitting the required inputs to model the quadruplex structure, the output interface on the right displays a consolidated summary of these inputs, and an interactive view of the generated model using JSmol. A video tutorial that succinctly explains the usage of the tool can be found at: https://youtu.be/U0VhdGoxP_I.

Through the HTML interface (**Figure 5**), the user has to provide information about the type of quadruplex motif (by selecting the appropriate image of the motif) which is to be used as a template for the fold, sequence type (DNA or RNA), number of quartets in the quadruplex fold, input residues individually for the 4 quadruplex strands (representing the G-quartets as well as non-G-quartets, if any), loop sequence as applicable, 5’ and 3’ overhang residues for each sequence (if applicable) and position of the non-G residues as in each strand. Note that in the case of a monomer, the sequence type should either be DNA or RNA, since the four strands of the quadruplex are connected. However, for the dimer (4 combinations), trimer (8 combinations), and tetramer (16 combinations), the sequence can be either DNA, RNA, or a combination of both. Soon after retrieving the information, the HTML interface passes it to a PHP script. The PHP subsequently decodes the quadruplex motif type into a 10-letter alphanumeric pattern. For instance, 1a-2p|3a-4 represents an antiparallel dimer motif named DA7 (**Figure S2**). Here, the first two strands (indicated by ‘1’ and ‘2’) are oriented in an antiparallel fashion (indicated by ‘a’), second and third (indicated by ‘3’) are oriented in a parallel fashion (indicated by ‘p’) and the third and fourth (indicated by ‘4’) strands are oriented in an antiparallel fashion (indicated by ‘a’). While ‘|’ indicates the separation of two independent sequences, ‘-’ indicates the two connected strands. Thus, the fold 1a-2p|3a-4 represents a dimer, wherein the strands 1 and 2 are formed by one sequence, and another sequence forms strands 3 and 4. Similarly, 1p-4p|3p-2 is a dimer (DP5) with a different connectivity. 1p-2p-3p-4 encodes for a monomer (named as MP1), wherein all four strands are connected in a parallel fashion. 1a|2a|3a|4 represents a tetrameric motif, QA3, in which the strands are in an antiparallel fashion with respect to each other. PHP passes this information to a bash script, which uses a FORTRAN code to generate the quadruplex fold coordinates and arranges the residue numbers as per the connectivity. Soon after the job is submitted, the user is provided with a molecule ID to retrieve the results later (**Figure 5**).

### Quadruplex fold generation and optimization

As mentioned above, using the appropriate helical twist and rise, the quadruplex folds having G and non-G quartets are generated. For the propeller loop, lateral loop, diagonal loop, diagonal-parallel loop, 5’-overhang, and 3’-overhang, the representative coordinates are initially generated as per the nucleotide sequence by applying helical rise using the bash script. Thus, the bash script generates the initial coordinates for the quadruplex folds, with steric hindrances and a disjointed backbone. Although the diagonal-parallel loop (*viz*., two alternate strands are connected in a parallel orientation) is not yet observed in any of the structures reported experimentally, the tool also models motifs having diagonal-parallel loops, since it is theoretically possible. Yet another point is that two crossing diagonals, which are present on the same side (DA3 motif in **Figure S2**), are also not observed experimentally but can be modeled by 3D-NuS-Qplex. In these cases, the user must consider a longer loop sequence to avoid steric hindrances. Subsequently, the generated models are subjected to the Powel minimization algorithm using Xplor-NIH [29], an updated version of Xplor [30], to establish backbone connectivity and remove steric hindrance.

Initially, 1000 cycles of Powell minimization are carried out by constraining (*viz*., not allowing them to move) the base atoms, leaving the backbone atoms free. During this round, only the dihedral energy terms are considered. For the subsequent rounds of minimization, quartet planarity and hydrogen bonding distance (using soft asymptotic potential) restraints, along with chi and sugar pucker restraints, are introduced. All the bonded energy terms are included, while van der Waals and electrostatic energy terms are included as discussed below. In the next step, the loop residues are kept free to remove the steric hindrance and establish the connectivity between adjacent nucleotides. If more than one loop is present (note that monomer has the highest number of 3 loops and tetramer has 0 loops), the loops are minimized one at a time. During this minimization cycle, all the quadruplex strands are also fixed. The minimization of a single loop occurs in six rounds, where each round involves 1,000 cycles of Powell minimization. During the first four rounds of minimization, atom-based nonbonded interactions (between the atoms whose mass is greater than zero) were considered with a cut-off value of 100 and by excluding the nonbonded interactions between bonded atoms, atoms that are bonded to a common third atom, and the atoms that are connected to each other through three bonds. While the electrostatic energy term is set to be off, the weightage of the van der Waals energy term is increased in each round (*viz*., 0.002, 0.005, 0.01, and 0.5). At the fifth round, the electrostatic energy is included, and the van der Waals energy term is given a weightage of 1. Atom-based nonbonded interactions are considered with a cut-off value of 4.5, and the non-bonded interactions are excluded as mentioned above, but with the inclusion of 1-4 nonbonded interactions. During the last round, 1000 cycles for Powell minimization are extended by increasing the weightage of the dihedral restraints to 2. Similarly, overhang residues are also minimized. Soon after all the loop residues are individually minimized (while constraining other loop and quadruplex strands), all the loop residues are left free for the next round of minimization. During the next round, each strand of the quadruplex is minimized individually to remove steric hindrance and establish internucleotide connectivity following the abovementioned six rounds of minimization protocol. Here, initially, the last (fourth) strand is left free while constraining the first three strands. Subsequently, the 3^rd^, 2^nd^, and 1^st^ strands are minimized one at a time. Following this, the fourth strand is minimized again as mentioned above, and with the inclusion of the 3^rd^, 2^nd^, and 1^st^ strands in subsequent steps. In the final round, all strands, loops, and overhang residues are not constrained and are only restrained; six rounds of minimization are carried out as mentioned above.

The user can access the results using the molecule ID provided during submission. Finally, the user can interactively view the modeled quadruplex fold with JSmol enabled within the HTML interface (**Figure 5**). The user can also download the PDB and XPLOR input files. A PHP script controls the entire process. **Figure 5** illustrates the overall workflow of 3D-NuS-Qplex, and **Figures S5** and **S6** provide representative examples of quadruplex models generated by 3D-NuS-Qplex.

## Conclusions

This study mathematically reveals 156 unique theoretically possible quadruplex motifs (two-layered quartets that act as the scaffold for the multi-layered folds) based on the strand connectivity and their relative orientations. While only 33 of these motifs have been found experimentally, the revelation of 156 possible motifs provides a new direction for exploring the diverse biological roles of these quadruplex folds. As learnt from the experimental structures, the combination of these motifs can further form complex higher-order quadruplex structures. Thus, one can envision that a greater number of higher-order quadruplex structures (arising from these 156 motifs) may exist in nature than those observed experimentally, adding further complexity to our understanding of these structures and their biological relevance. A web tool, 3D-NuS-Qplex, is also developed here to model all the theoretically possible quadruplex folds, incorporating 5’ and/or 3’-overhangs, non-G-quartets, and different combinations of DNA and/or RNA strands. Since the quadruplex structures are found to play a crucial role in several biological functions and diseases, this study brings a new dimension to the (i) understanding of quadruplex fold dependency on biological functions and diseases, (ii) fold-dependent design of ligands, and (iii) custom design of synthetic quadruplex folds for nanotechnological applications.

## Author contribution

SS developed the theoretically possible quadruplex motifs, designed the HTML interface, and carried out testing. TR developed the methodology, implemented the modeling in PHP and bash scripting. SS and TR wrote the manuscript. TR designed and supervised the project.

## Acknowledgements

The authors thank Dr Abhishek Kumar for the initial discussion on the 3D-NuS-Qplex web tool and the IITH computer center for technical support. SS thanks the Ministry of Education (MoE) for the fellowship.

